# QNPtoVox: A methods pipeline for mapping 2D quantitative neuropathology to 3D MNI voxel space

**DOI:** 10.64898/2026.04.17.719274

**Authors:** Raghav Madan, Tejas Bajwa, Amanda Kirkland, David Hunt, Jason Webster, Seo-Eun Choi, Christine L. Mac Donald, Nadia Postupna, Caitlin S. Latimer, Thomas J. Grabowski, John H. Gennari, Paul K. Crane

**Affiliations:** Department of Biomedical Informatics and Medical Education, University of Washington, Seattle, WA, USA; Division of Neuropathology, Department of Laboratory Medicine and Pathology, University of Washington, Seattle, WA, USA; Department of Neurological Surgery, UW School of Medicine, Seattle, WA, USA; Department of Radiology, University of Washington, Seattle, WA, USA; School of Medicine, University of Washington, Seattle, WA, USA; Department of Neurology, University of Washington, Seattle, WA, USA

## Abstract

Quantitative neuropathology has advanced through whole-slide imaging and digital histology platforms. Yet, these measurements rarely align with neuroimaging coordinate frameworks that may be useful for spatial modeling and other applications. QNPtoVox, short for quantitative neuropathology to voxels, is a reproducible, modular pipeline that transforms quantitative metrics generated by digital pathology software (HALO) into voxel-based maps registered to a standard common coordinate (MNI) template. The workflow integrates digital histopathology, gross tissue photography, ex-vivo MRI, and nonlinear registration to generate spatially standardized 3D pathology representations. This Methods article provides a complete procedural description, including required materials, step-wise instructions, operator-dependent checkpoints, expected outputs, reproducibility evaluation, and troubleshooting. QNPtoVox enables voxel-level integration of neuropathology with neuroimaging tools, unlocking existing histopathology datasets for computational modeling and cross-cohort harmonization.

## 2. Introduction

Quantitative neuropathology has undergone a rapid transformation over the past decade, driven by advances in whole-slide imaging^1^, high-throughput digital histology, and computational analysis platforms. Modern image analysis tools, including deep learning classifiers and quantitative immunohistochemical pipelines, now enable digitization and analysis of brain tissue sections at subcellular resolution. Software systems such as Halo^2^, QuPath^3^, and other pixel-level analytical environments have enabled objective quantification of pathological markers, including tau, amyloid, α-synuclein, TDP-43, and microvascular injury, across entire tissue sections. These technologies have ushered in an era in which neuropathological features can be measured in three-dimensional, continuous space rather than through coarse, region-level scoring schemes such as Braak staging^4,5^, CERAD^6^, or Thal phases^7,8^.

Despite the unprecedented richness of modern digital histopathology, quantitative neuropathology remains fundamentally limited by its spatial isolation. Measurements are taken in a two-dimensional histological plane, divorced from the brain’s three-dimensional architecture. As a result, quantitative pathology is complex to integrate with spatially structured information derived from MRI-based modalities, including structural MRI, diffusion MRI, resting-state fMRI, and PET. This dividing line is becoming more problematic as the field advances toward computational models of neurodegeneration that explicitly depend on interconnected anatomical information. These models assume spatial continuity, but quantitative histopathology, in practice, lacks this critical alignment. Consequently, the most biologically detailed measurements (histology) remain analytically disconnected from the broader spatial context (neuroimaging) where brain-wide disease processes can be visualized.

Recent advances in ex vivo MRI provide an essential bridge between these domains. High-resolution MRI performed on fixed postmortem tissue preserves gross anatomical architecture, enabling direct correspondence between the brain’s macroscopic structure and the microscopic features observed on histology. Ex vivo MRI resolves cortical layers, subcortical nuclei, white matter tracts, and the geometry of grossly sectioned tissue slabs, and it captures morphological characteristics before tissue distortion occurs during histological processing. Researchers have increasingly recognized the potential of ex vivo MRI as an alignment scaffold for histological data, as demonstrated by Webster et al. (2021)^9^, who used ex vivo MRI to evaluate sampling precision in Alzheimer’s disease neuropathology protocols. Their work illustrates the feasibility of using MRI to reintroduce anatomical context into tissue-based measures. However, their approach, and others like it, stop short of providing a full computational pipeline for spatially registering quantitative pathology into a coordinate system compatible with neuroimaging tools.

A second technological advance that makes the work described here possible is the growth of standardized brain coordinate systems, most prominently the Montreal Neuroimaging Institute’s (MNI) ICBM 2009^10^ series. These templates encode a population-representative volumetric space with consistent orientation, dimensions, and anatomical landmarks. The MNI framework anchors nearly all modern neuroimaging tools, FreeSurfer^11^, FSL^12^, SPM^13^, and ANTs^14^. It also supports anatomical parcellations, connectivity atlases, tissue probability maps, white matter tractography, and functional networks. Hundreds of group-level resources exist in MNI space, enabling cross-study harmonization and comparative neuroanatomy. Yet, quantitative neuropathology has not been brought into this shared coordinate space, leaving a deep gap between histology and neuroimaging.

QNPtoVox fills this critical methodological gap. It is a modular, lightweight, and easy-to-implement pipeline that maps Halo^2^-derived quantitative pathology tiles into voxel-level maps in MNI space through a series of explicit computational steps. The method requires no specialized hardware beyond standard histological workflows and uses transparent, script-based transformations that can be run on a workstation. Most importantly, the pipeline makes existing neuropathology datasets spatially analyzable, unlocking decades of histopathological material for integration with MRI-anchored computational frameworks.

In this article, we describe QNPtoVox in detail, including materials, software requirements, step-by-step procedures, validation, operator checkpoints, anticipated results, and troubleshooting. Our goal is to equip neuropathology and neuroimaging laboratories with a reproducible workflow for generating spatially standardized, voxel-level pathology maps, thereby bridging the gap between histological precision and the anatomical context of neuroimaging.

## 3. Materials and equipment

We derived all biological materials and imaging data used in the QNPtoVox pipeline from standardized neuropathological procedures described in detail by Latimer et al. (2023)^15^. The protocol defines systematic fixation, regionally targeted tissue sampling, immunohistochemistry, and both qualitative and quantitative assessment of postmortem human brain tissue. In developing QNPtoVox, we followed this established tissue-handling and processing framework, including formalin fixation, ex vivo MRI acquisition, coronal slab preparation, region-specific tissue sampling, and AT8 immunohistochemistry for phosphorylated tau (clone AT8, Thermofisher, MN1020, 1:1000 dilution). By adhering to these validated methodological standards, we ensured consistency, reproducibility, and high-quality inputs for downstream digital pathology and MRI-based spatial processing.

We extracted brains using standard postmortem procedures and immersion-fixed them in formalin. After fixation, we acquired ex vivo MRI scans of the fixed tissue through the Adult Changes in Thought^16^ (ACT) study’s Neuropathology and Neuroimaging Cores, following the standardized postmortem imaging workflow described by Latimer et al. (2023). We scanned each fixed hemisphere on a 3T system using high-resolution FLAIR sequences at approximately 800 µm isotropic resolution. Within QNPtoVox, these ex vivo MRI volumes serve as the primary three-dimensional anatomical scaffold for downstream spatial alignment.

Following MRI acquisition, we sectioned the brains coronally and photographed each tissue slab at the time of dissection using calibrated, ruler-referenced gross imaging. We use these gross photographs solely to verify slice-level correspondence between histological sections and the ex vivo MRI volumes. Although we do not perform quantitative registration on these images, they provide essential anatomical context during spatial alignment and quality control.

From the coronal slabs, we sampled region-specific tissue blocks according to standardized neuropathological protocols^17^. We processed these tissue blocks for paraffin embedding, sectioning, and immunohistochemical staining using the AT8 antibody to label phosphorylated tau. We then generated digital whole-slide images from the AT8-stained sections using high-resolution brightfield scanners operated by the University of Washington Neuropathology Core. We further performed quantitative digital pathology using Halo^2^, and these measurements constitute the primary histopathological inputs to the QNPtoVox pipeline. Because this manuscript focuses on computational spatial registration, we do not describe laboratory processing, staining, or scanning procedures in detail here; instead, we refer readers to Latimer et al. (2023)^15^ for the complete experimental protocol.

### 3.1 Participants and Inclusion Criteria

We developed QNPtoVox using data from 10 participants in the ACT Study^16^, a population-based longitudinal cohort of older adults followed until autopsy. We selected participants based on the availability of complete data across all three modalities required by the QNPtoVox workflow: high-quality AT8 whole-slide images suitable for quantitative analysis, matched gross coronal slab photographs, and ex vivo MRI acquired before tissue sectioning. We included only participants with intact middle frontal gyrus (MFG) slabs and successful Halo-based quantitative pathology processing. We excluded cases with excessive tissue damage, poor staining quality, or incomplete MRI coverage. To ensure sufficient tau burden for spatial modeling, we required all included participants to meet Braak stages IV-VI criteria, reflecting moderate to severe tau pathology.

### 3.2 Computational environment and software dependencies

We implemented the QNPtoVox pipeline using a combination of open-source software tools and widely used neuroimaging platforms. We obtain quantitative histopathology measurements using Halo^2^, which we use exclusively for the initial extraction of tile-level pathology values and spatial annotations. We perform all subsequent processing steps using publicly available, platform-independent, script-based tools.

We implemented pipeline orchestration and coordinate processing in Python^18^ and relied on standard scientific libraries, including NumPy^19^, pandas^20^, SciPy^21^, and NiBabel^22^, for numerical computation, data handling, and volumetric image processing. These libraries support voxel-wise data extraction and operate directly on NIfTI-formatted images throughout the pipeline. We implemented virtual slicing of ex vivo MRI volumes in R^23^ using the RNifti^24^ package for NIfTI input–output and the PNG^25^ package for slice-level image export. We adapted the slicing script from previously published virtual pathology tools and incorporated it into the QNPtoVox codebase.

We rely on established neuroimaging software packages for conversion and preprocessing of neuroimaging formats. We use FreeSurfer^11^ to convert postmortem MRI volumes from MGZ to NIfTI format. We perform manual reorientation and upsampling of MRI volumes, along with visual quality control of intermediate outputs, using FSL^12^ and FSLeyes^26^. We perform nonlinear registration of participant-specific volumes to the MNI ICBM 2009b^27^ template using the Advanced Normalization Tools (ANTs) software suite^14^.

We developed and tested the QNPtoVox pipeline on Unix-based operating systems, including Linux and macOS. We designed the pipeline to run without specialized hardware or graphical processing units, and we typically complete runs on a standard workstation with sufficient memory to accommodate high-resolution MRI volumes. We provide detailed installation instructions, software version recommendations, and guidance on environment setup in the accompanying GitHub repository.

**GitHub repository:** https://github.com/RaghavMadan/QNPtoVox

## 4. Methods

### 4.1 Overview of the computational workflow

The computational workflow described here transforms two-dimensional quantitative neuropathology measurements into three-dimensional, voxel-aligned representations in MNI standard neuroimaging coordinate space. To support clarity and reproducibility, we divide the workflow into two conceptual steps.

**Step 1** focuses on extracting quantitative pathology and associated spatial coordinates from whole-slide histology using Halo. This step produces tile-wise measurements in native histological coordinate space.

**Step 2** encompasses the full QNPtoVox pipeline. This stage integrates histological coordinates with ex vivo MRI anatomy, performs slice-level and volumetric alignment, and generates voxel-based pathology maps that can be registered to MNI space for downstream analysis.

### 4.2 Step 1- Quantitative neuropathology extraction using Halo

In the first stage of the workflow, we extract quantitative neuropathology measurements and their associated two-dimensional spatial coordinates from digitized histological sections using Halo^2^. This step generates the foundational quantitative and geometric inputs required for all downstream spatial transformations performed by the QNPtoVox pipeline. T.B. developed and implemented this initial extraction step.

We load AT8-immunostained whole-slide images into Halo following standard digital pathology preprocessing and quality control procedures established by the ACT Neuropathology Core^15^. We developed the pipeline using tissue from the cortical grey matter and adjacent white matter of the middle frontal gyrus. For each section, we manually delineate cortical regions of interest using freehand annotations to exclude artifacts, tissue folds, and non-cortical structures. We favor manual annotation over automated tissue detection to ensure precise exclusion of spurious regions and to preserve anatomical fidelity for downstream spatial modeling.

After defining cortical regions of interest, we overlay a uniform 2D grid with a 0.5 × 0.5 mm spacing on each annotated tissue section. We selected this grid resolution to approximate the 0.5 mm isotropic voxel size of the MNI ICBM 2009b template, which is used later in the pipeline, thereby facilitating conceptual alignment between histological sampling units and voxel-based representations. We retain only grid tiles that fully meet the specified dimensions and exclude partial tiles at tissue boundaries to maintain consistency of spatial units.

**Figure 1:**
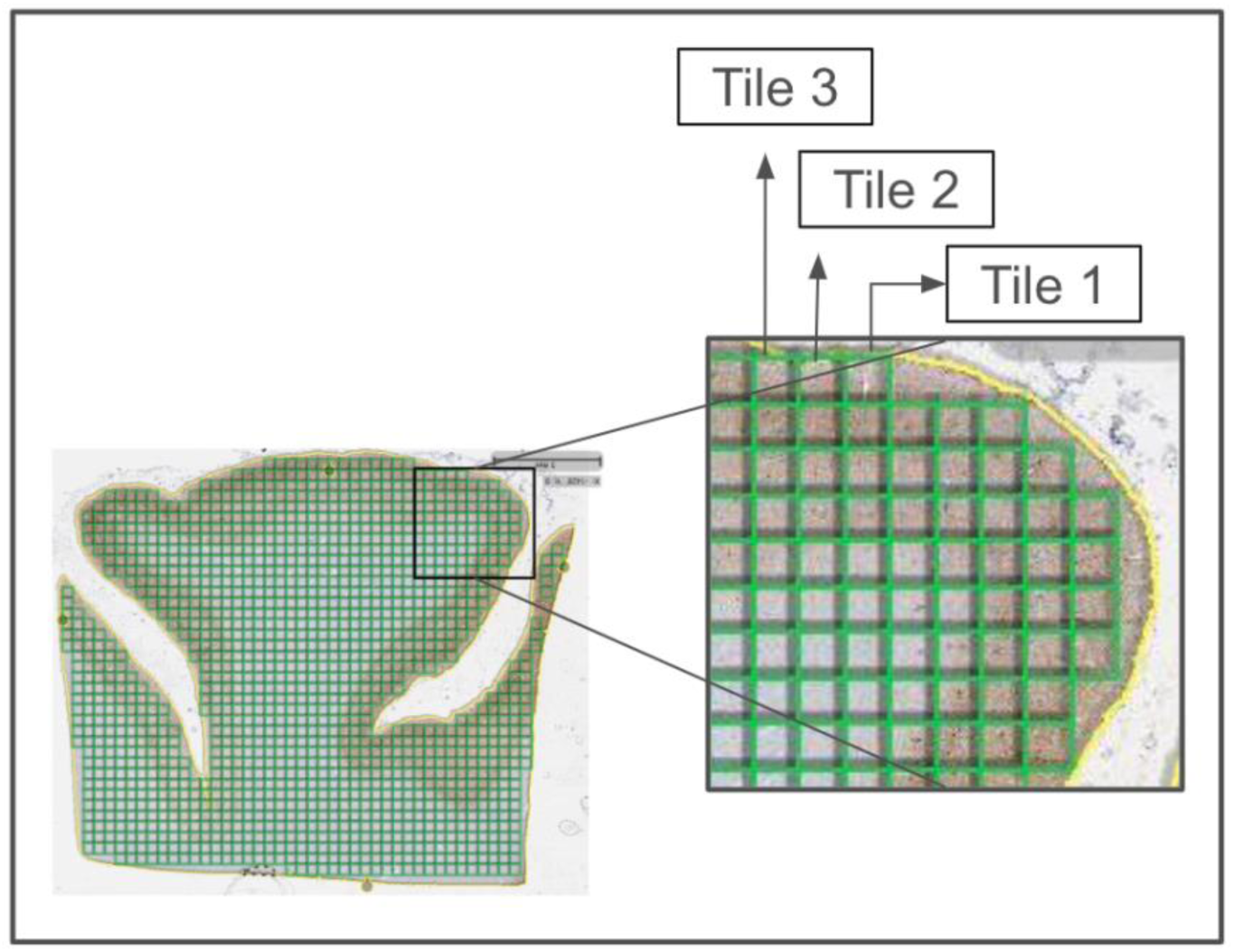
A uniform two-dimensional grid with 0.5 × 0.5 mm spacing is overlaid on a manually delineated AT8-stained histological section, encompassing cortical grey matter and adjacent white matter in the middle frontal gyrus region.

We calculate quantitative tau pathology within each grid tile using Halo’s Area quantification module. The algorithm identifies AT8-positive signal using a color-based classification approach, and we adjust stain-specific parameters to account for staining intensity variability across slides. For each grid tile, Halo computes the percent area positive for AT8 staining, yielding a continuous quantitative measure of tau burden. We perform quality control by visually inspecting algorithm outputs and adjusting optical density parameters accordingly.

In parallel with quantitative analysis, we export spatial annotation data from Halo to preserve the relative geometry of the grid tiles. We represent each tile as a closed square in the two-dimensional coordinate space of the histological image, with vertex coordinates recorded in microns. This representation preserves the spatial arrangement of tiles across the tissue section while remaining agnostic to three-dimensional anatomical orientation.

In this step, we generate two primary data products. First, we export a summary table containing quantitative pathology values, including the percent area positive for AT8 staining for each grid tile. Second, we export an XML annotation file containing the spatial coordinates for each grid tile. Together, these files encode both the magnitude of pathology and the spatial relationships among sampling units in histological space.

These outputs constitute the complete input to the QNPtoVox spatial pipeline. At this stage, the extracted quantitative values and two-dimensional coordinates are not yet aligned to brain anatomy or template space. However, they preserve all information required for subsequent co-location with ex vivo MRI, voxel transformation, and registration to MNI space. We can store the exported Halo annotation and summary files indefinitely and reuse them for repeated pipeline execution or sensitivity analyses without rerunning histological quantification.

### 4.3 Step 2- QNPtoVox: Spatial transformation of quantitative neuropathology into voxel and MNI space

In Step 2, we transform tile-level quantitative neuropathology data into three-dimensional voxel representations aligned to the MNI space. This step operationalizes the spatial integration of histology and imaging by orchestrating pipeline execution, preprocessing ex vivo MRI volumes, and enforcing standardized input–output dependencies. The following subsections describe the execution framework and the preprocessing steps that establish the anatomical scaffold for downstream co-location, voxelization, and registration to MNI space. R.M. developed and implemented QNPtoVox.

#### 4.3.1 Pipeline orchestration and execution framework

We control pipeline execution through a central Python script (*run_qnp_pipeline.py*), which serves as the main entry point for running one or more processing stages across selected participants. We designed the pipeline to be modular, allowing us to execute individual steps independently or chain them together to support iterative development, manual quality control, and batch processing. Configuration files specify participant lists, directory paths, and step-specific parameters, enabling us to reuse the same codebase across datasets without modification.

Each pipeline stage follows a standardized input–output convention and writes participant-specific outputs to a structured directory hierarchy. The pipeline automatically logs execution status, warnings, and errors, ensuring full traceability of all processing steps.

#### 4.3.2 Preprocessing of ex vivo MRI volumes

To establish an anatomical scaffold for spatial alignment, we first standardized each participant’s ex vivo MRI volume in terms of resolution, orientation, and format. Input MRI data consist of postmortem scans acquired from fixed hemispheres and stored in FreeSurfer’s^11^ MGZ format. We manually convert these volumes to NIfTI format, reorient them to match the MNI 2009b anatomical axes, and upsample them to 0.5 mm isotropic resolution using FSLeyes^26^. We perform this manual reorientation and upsampling step to accommodate inter-participant variability in tissue geometry and to ensure accurate correspondence with template space. Specific information about these steps is available in the FSLeyes^26^ software documentation.

The resulting upsampled and reoriented MRI volume serves as the anatomical reference for all downstream slicing, co-location, and voxelization operations. We halt pipeline execution if this prerequisite file is missing, thereby enforcing strict dependency management.

**Figure 2:**
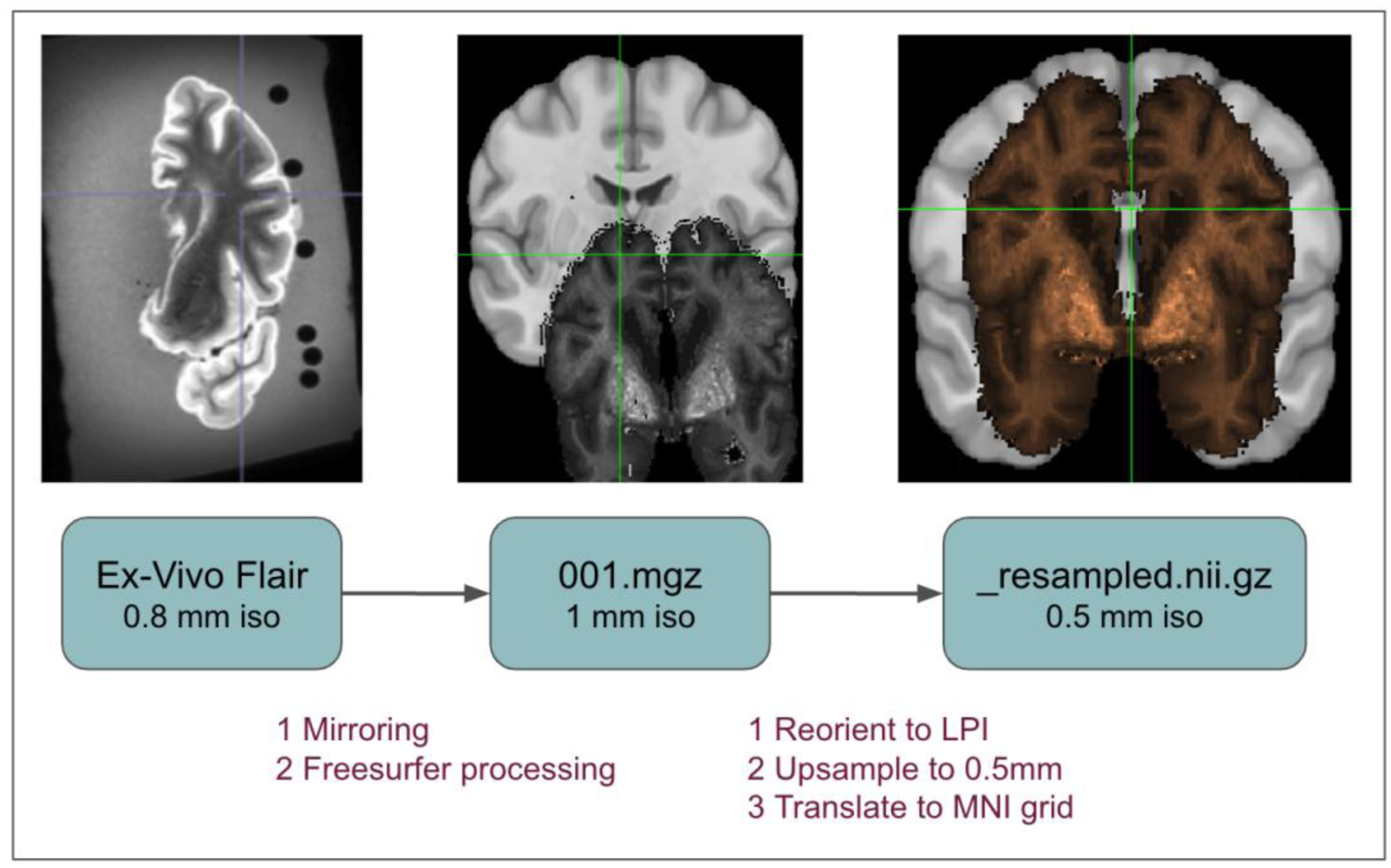
Preprocessing workflow for ex vivo MRI volumes. Starting with a participant’s original FLAIR scan (left), the image is mirrored and converted to FreeSurfer’s^11^ .mgz format (center), then reoriented to LPI (Left-to-Right, Posterior-to-Anterior, Inferior-to-Superior), upsampled to 0.5 mm isotropic resolution, and translated to the MNI grid (right). The final output serves as the anatomical reference for voxel-level integration with QNP data.

#### 4.3.3 Virtual slicing of ex vivo MRI

To establish a slice-wise anatomical reference system, we virtually slice the upsampled ex vivo MRI volume into a series of coronal images at 0.5 mm intervals. We implement this step using a custom R script adapted from the virtual Pathology toolkit^9^ and integrated into the QNPtoVox pipeline execution framework.

The slicing procedure automatically identifies the anterior and posterior extents of brain tissue by detecting non-zero voxel intensities along the coronal axis. By doing so, we exclude empty padding while retaining all tissue-containing planes. We intensity-normalize each coronal slice and reorient it into a standardized anatomical view to ensure consistency with gross tissue photographs and Halo-derived histology images. We save the resulting slice images as PNG files using a naming convention that preserves the original slice index, thereby enabling direct traceability between the two-dimensional images and the three-dimensional MRI volume. These virtual coronal slices provide a standardized visual substrate for expert-guided anatomical correspondence in the subsequent co-location step.

#### 4.3.4 Expert-guided co-location of histopathology with ex vivo MRI

We achieve accurate spatial alignment between histological samples and MRI anatomy through an expert-guided co-location step. For each participant, a trained neuroanatomist (A.K.) identifies the coronal MRI slice that corresponds to the physical tissue slab and determines the in-plane anatomical location of the sampled tissue within that slice.

To make this determination, we perform a systematic visual comparison of three data sources: the virtual coronal MRI slices generated in the previous step, gross photographs of the coronal tissue slabs acquired during dissection, and Halo-derived digital histology images corresponding to the sampled tissue block. The neuroanatomist uses macroscopic anatomical landmarks, including sulcal and gyral patterns, cortical curvature, ventricular configuration, and slab geometry, to identify the relevant coronal plane. The neuroanatomist then uses finer-scale anatomical features visible in histological and gross images to localize the sampling region within that plane.

We record the identified coronal slice index and the estimated in-plane offsets in a participant-specific configuration file. These values serve as spatial anchors for mapping two-dimensional histological tile coordinates into three-dimensional MRI voxel space. By externalizing these expert decisions into configuration files rather than hard-coding them, we preserve transparency, reproducibility, and auditability within the QNPtoVox pipeline.

**Figure 3:**
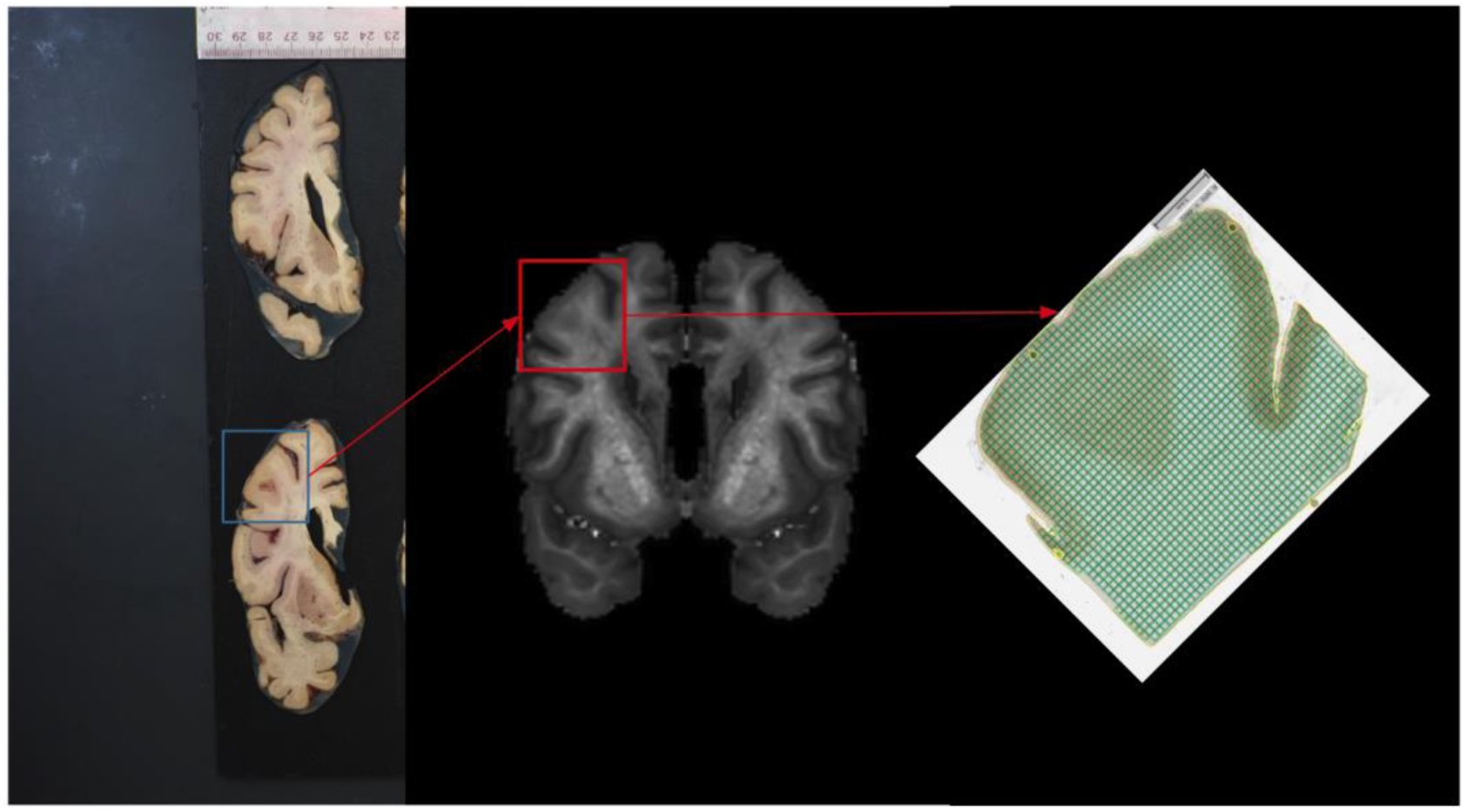
Co-location of quantitative neuropathology with ex vivo MRI for participant 7038. The left panel shows the gross tissue slab used for histological sampling. The middle panel displays a coronal slice from the participant’s upsampled ex vivo MRI volume. The right panel shows the corresponding Halo-annotated tissue section overlaid with a 0.5 × 0.5 mm tile grid. Red arrows illustrate the anatomical alignment process used to identify the Y-plane and apply coordinate offsets for accurate spatial mapping.

#### 4.3.5 Mapping histological coordinates into voxel space

Using the spatial anchors defined during the expert-guided co-location step, we automatically transform Halo-derived tile coordinates from two-dimensional histological space into three-dimensional voxel coordinates within each participant’s native MRI volume. Once the coronal slice index and in-plane offsets are specified, the pipeline executes this transformation without further manual intervention.

For each tile, we compute the centroid in histological coordinate space, place it at the manually identified coronal plane, and shift it according to the recorded in-plane offsets. This procedure preserves the relative spatial arrangement of pathology across the tissue section while embedding the data within the three-dimensional anatomical context of the MRI volume.

The resulting transformed coordinates serve as the basis for constructing volumetric representations of pathology. We expand each histological tile into a three-dimensional block corresponding to the physical thickness of the sampled tissue, thereby generating an initial voxel-wise pathology mask in native MRI space. This automated voxelization step produces a structured volumetric representation that is subsequently refined through alignment and smoothing operations.

**Figure 4:**
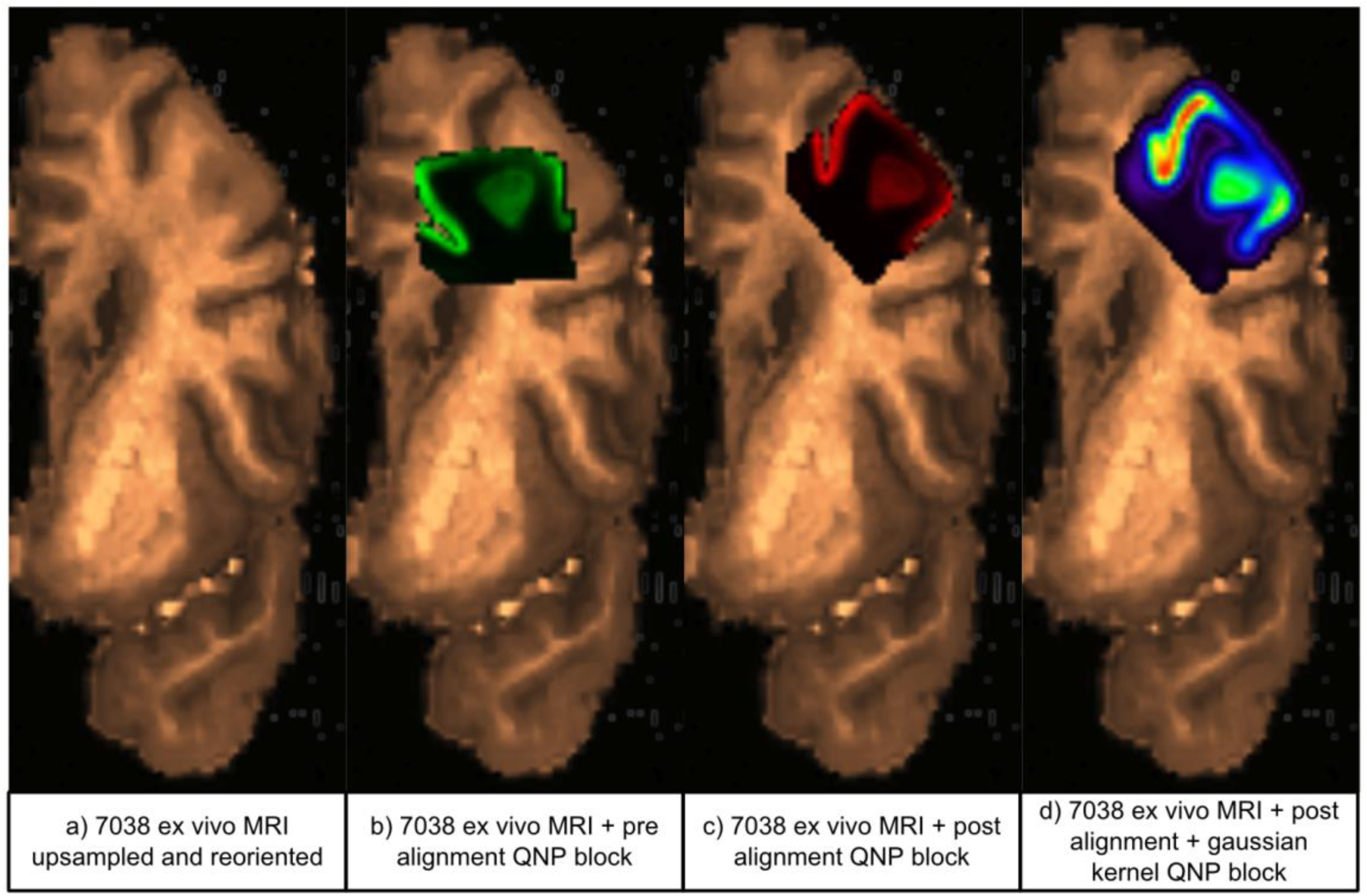
Transformation of QNP block to voxel space for participant 7038. (a) Upsampled and reoriented ex vivo MRI in native space. (b) Initial placement of the QNP block based on transformed coordinates prior to alignment. (c) Manually aligned QNP block adjusted to match anatomical boundaries using FSLeyes^26^. (d) Final smoothed pathology map after application of a 2 mm Gaussian kernel, illustrating local diffusion and anatomical conformity.

#### 4.3.6 Volumetric smoothing and preparation for registration

To account for spatial uncertainty in tissue sampling and to generate biologically plausible pathology maps, we smooth the voxelized pathology volume using a three-dimensional Gaussian kernel. This operation locally distributes pathology signal, mitigates hard-edged artifacts introduced during block construction, and improves robustness for downstream voxel-wise analyses. The resulting smoothed volume provides a continuous estimate of pathology density within each participant’s brain.

#### 4.3.7 Registration to the MNI space

In the final stage of the QNPtoVox pipeline, we register participant-specific pathology volumes to the MNI ICBM 2009b nonlinear symmetric template. We estimate nonlinear registration parameters by aligning each participant’s upsampled ex vivo MRI volume to the template using established deformable registration tools (full implementation details available at the GitHub repository). We then apply these transformations to the smoothed pathology volume, yielding a voxel-aligned pathology map in standard space.

The final output of the pipeline is a three-dimensional pathology volume in MNI space that we can directly integrate with neuroimaging atlases, voxel-wise covariates, and spatial modeling frameworks. This standardized representation enables cross-participant comparison, group-level analyses, and integration with multimodal neuroimaging data.

**Figure 5:**
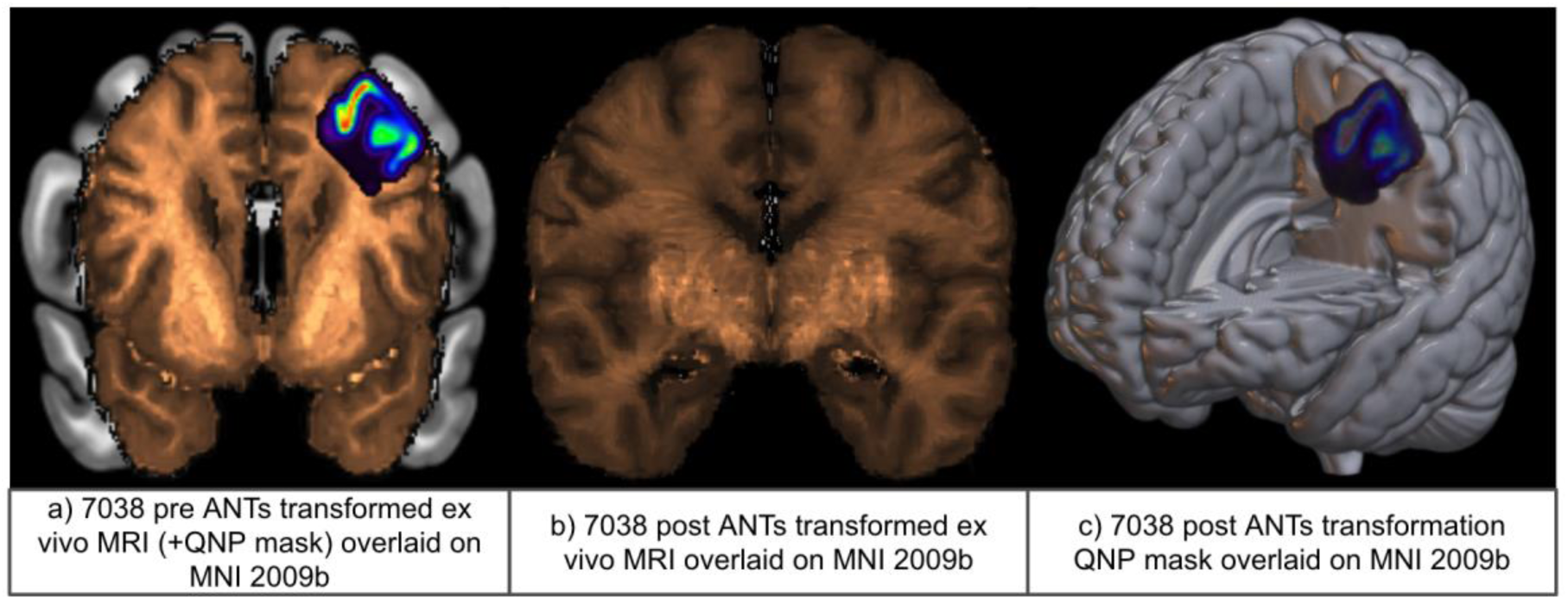
Registration of participant 7038’s ex vivo MRI and QNP volume to MNI space using ANTs^14^ registration tool. (a) Pre-ANTs transformation view showing the native-space ex vivo MRI overlaid with the aligned QNP mask. (b) Post-ANTs transformation of the ex vivo MRI, nonlinearly warped and aligned to the MNI2009b template. (c) Final smoothed QNP mask registered to MNI space and visualized on the MNI2009b cortical surface.

#### 4.3.8 Voxel-data extraction

Although QNPtoVox operates primarily on volumetric data during transformation and registration, we define the final analytical product as a voxel-coordinate database rather than a standalone image file. After registering pathology volumes to MNI space, we programmatically parse the resulting volume to identify all voxels containing non-zero signal. For each such voxel, we extract its spatial coordinates and associated pathology intensity and write these values to a structured tabular file.

This final extraction step ensures that QNPtoVox outputs are directly usable in computational workflows that operate on structured data rather than image volumes. By decoupling spatial representation from analytical method, we maximize reuse and interoperability with existing modeling and spatial analysis frameworks.

#### 4.3.9 Data and code availability

The QNPtoVox pipeline is implemented as a modular, script-based workflow and is publicly available to support transparency, reproducibility, and reuse. All code required to execute the pipeline, including pipeline orchestration scripts, processing modules, configuration templates, and documentation, is hosted in a publicly accessible GitHub repository.

The repository includes step-by-step instructions for environment setup, pipeline execution, and manual intervention steps, as well as example configuration files that illustrate expected directory structures and parameter settings. Source code for all pipeline components, including MRI preprocessing, virtual slicing, histology–MRI co-location, voxel transformation, smoothing, and MNI registration, is provided in full.

Due to participant privacy and data-use agreements, the raw histological images, gross tissue photographs, and ex vivo MRI data used in this study are not publicly available. These data were obtained from the Adult Changes in Thought (ACT) study and are subject to controlled access. Researchers interested in accessing the underlying biological or imaging data should contact the appropriate data governance bodies associated with the ACT study.

All derived outputs generated by the QNPtoVox pipeline, including intermediate volumes and voxel-coordinate pathology datasets, can be reproduced by running the pipeline on authorized input data using the provided code and documentation. No proprietary software is required beyond the digital pathology platform used for initial histological quantification.

## 5. (Anticipated) Results

Successful execution of the QNPtoVox pipeline produces a spatially standardized, voxel-coordinate–based representation of quantitative neuropathology for each participant. The central outcome of the method is a dataset in which quantitative pathology measurements derived from digital histology are expressed at the level of individual voxels within a common three-dimensional neuroanatomical coordinate system. This transformation enables quantitative neuropathology to be analyzed using the same spatial frameworks routinely applied to neuroimaging data.

At the conclusion of the pipeline, each participant is associated with a three-dimensional pathology volume registered to the MNI ICBM 2009b nonlinear symmetric template. In this volume, voxel intensities represent continuous estimates of tau pathology burden derived from Halo-quantified percent area stained for AT8, spatially distributed via voxelization and smoothing. Because all volumes are aligned to a common template space, corresponding voxels across participants refer to homologous anatomical locations, enabling direct comparisons and aggregations across individuals.

In addition to the final MNI-aligned output, the QNPtoVox pipeline generates a series of intermediate volumetric and slice-level outputs that support visual inspection and quality control. At each central stage – upsampled native-space MRI, coronal slice generation, initial pathology block placement, post-alignment pathology volumes, and smoothed representations – users can review intermediate files to confirm anatomical consistency. When the pipeline is functioning as intended, these interim volumes exhibit apparent anatomical coherence between histopathological sampling and brain anatomy. Grey and white matter boundaries in the pathology volumes align with those visible in the ex vivo MRI, cortical ribbon thickness is preserved, and major gyral and sulcal features overlap across modalities. At the slice level, the spatial extent and orientation of tissue blocks closely match the underlying cortical geometry in the MRI, providing intuitive confirmation of correct co-registration.

To facilitate downstream statistical analysis and modeling, the volumetric pathology representation is subsequently converted into a structured, voxel-coordinate database. After registering to the MNI space, the pathology volume is parsed to identify all voxels with non-zero signal. For each such voxel, the three-dimensional MNI coordinates and the associated quantitative pathology value are extracted and recorded. The resulting dataset consists of rows representing individual voxels, each defined by its spatial coordinates in template space and a continuous pathology measurement, with participant identifiers retained to support both within- and across-subject analyses.

This voxel-coordinate representation constitutes the primary analytical output of QNPtoVox. By expressing quantitative neuropathology in tabular form rather than solely as an image volume, the pipeline enables direct integration with statistical and computational modeling frameworks that operate on structured data. At the same time, because the data remain anchored to MNI coordinates, they can be readily aggregated into atlas-defined regions, projected onto cortical surfaces, or combined with voxel-wise neuroimaging covariates as required by the analytical context.

The outputs exhibit several defining properties that distinguish them from traditional neuropathological representations. Pathology values are continuous rather than ordinal or categorical, preserving the full dynamic range of quantitative histology. Spatial resolution is defined at the voxel level rather than predefined regions of interest, allowing flexible post hoc modeling strategies. Importantly, each voxel-level observation remains traceable to its histological origin through the pipeline’s intermediate outputs, supporting interpretability, debugging, and reproducibility.

When the pipeline executes successfully, visual inspection of both intermediate and final outputs should demonstrate anatomically plausible localization of pathology within the targeted cortical region, with consistent alignment across participants in template space. The resulting voxel-coordinate database provides a reusable, extensible foundation for spatial modeling, multimodal integration, and hypothesis-driven analysis, and serves as the key deliverable of the QNPtoVox method.

Although QNPtoVox provides a reproducible framework for mapping quantitative neuropathology into voxel and template space, several limitations and practical considerations should be acknowledged. Many of these arise from the inherent complexity of linking two-dimensional histology with three-dimensional brain anatomy and reflect trade-offs common to multimodal postmortem workflows.

A primary limitation of the pipeline is its reliance on expert-guided anatomical co-location. Identification of the coronal MRI slice corresponding to a given histological tissue block and estimation of in-plane offsets require neuroanatomical expertise and visual judgment. Although this step is constrained by standardized virtual slicing, gross tissue photography, and consistent anatomical landmarks, some degree of operator variability is unavoidable. To mitigate this, QNPtoVox externalizes all manually determined parameters into configuration files and preserves intermediate outputs at each stage, allowing decisions to be reviewed, revised, or replicated by independent operators. Users are encouraged to document the rationale for slice selection and to perform visual cross-checks using multiple anatomical cues.

The accuracy of spatial mapping is also dependent on the quality and completeness of the ex vivo MRI data. Distortions introduced during fixation, embedding, or scanning can affect the fidelity of cortical boundaries and sulcal geometry, particularly in regions with substantial postmortem deformation. When tissue damage or imaging artifacts obscure anatomical landmarks, co-location may be less precise. Visual inspection of intermediate MRI volumes and slice-level outputs is therefore a critical quality control step, and users should verify that grey and white matter boundaries, cortical ribbon thickness, and major gyral features are preserved prior to proceeding with voxelization and registration.

Another limitation relates to the spatial sparsity of histological sampling. QNPtoVox operates on pathology data derived from discrete tissue blocks rather than exhaustive whole-brain histology. As a result, the resulting pathology volumes represent spatially informed estimates rather than direct measurements at every voxel. The application of volumetric smoothing partially addresses this limitation by distributing the pathology signal locally. Still, users should interpret voxel-level values in the context of sampling density and kernel parameters. Kernel size and smoothing choices influence the spatial extent of inferred pathology and should be selected with consideration of the anatomical scale and scientific question of interest.

The current implementation of QNPtoVox has been validated within a single cortical region and pathology marker. Although the pipeline is extensible to additional regions, stains, and cohorts, generalization beyond the tested context may require adjustment of configuration parameters, alignment strategies, or quality control thresholds.

Common troubleshooting scenarios involve missing or misnamed input files, inconsistencies between configuration parameters and directory structure, or failures during stepwise execution due to unmet dependencies. The pipeline includes input validation and logging mechanisms to identify such issues early. When errors occur, users should consult the log files generated during execution and verify that all prerequisite steps have been completed successfully. In particular, failure to generate the upsampled MRI volume or to populate manual co-location parameters will prevent downstream steps from executing correctly.

Despite these limitations, QNPtoVox is designed to make sources of uncertainty explicit rather than implicit. By preserving intermediate volumes, logging all transformations, and separating automated processing from expert-guided decisions, the pipeline supports transparency, reproducibility, and iterative refinement. When used with appropriate quality control and anatomical oversight, it provides a robust and flexible framework for spatially integrating quantitative neuropathology with neuroimaging coordinate systems.

## 6. Discussion

We developed QNPtoVox to address a central methodological limitation in quantitative neuropathology: the absence of a shared anatomical coordinate framework that allows spatially explicit pathology measurements to be compared, aggregated, and modeled across individuals. A common reference space establishes voxel-level comparability, ensuring that a voxel at a given anatomical location, such as the left middle frontal gyrus, corresponds to the same spatial point across brains. Without such a framework, quantitative pathology measurements remain spatially isolated, limiting meaningful aggregation and cross-participant analysis. By aligning pathology to the MNI template, QNPtoVox enables direct integration with multimodal neuroimaging features, including structural MRI, diffusion-derived connectivity, vascular territories, tissue probability maps, and functional networks. This alignment supports anatomically grounded models of regional vulnerability and disease spread, and it situates quantitative histology within the many high-resolution anatomical and functional atlases that exist exclusively in template space. More broadly, standardized coordinate systems provide the three-dimensional coherence required for computational models of neurodegeneration and enable standardized, reusable, and benchmarkable analytic workflows across studies and research groups.

Importantly, spatial registration of quantitative neuropathology to each participant’s native brain anatomy is an inherent byproduct of the QNPtoVox pipeline. Before transformation to template space, QNPtoVox produces voxel-aligned pathology volumes in native ex vivo MRI space, preserving individual-specific anatomy and tissue geometry. These native-space representations can be used independently of MNI registration for subject-level analyses when cross-participant comparison is not required. For example, native-space quantitative neuropathological maps can support within-subject investigations of regional pathology gradients, comparisons between histology and subject-specific imaging features, or visualization of pathology burden relative to individual cortical folding patterns and lesion distributions. This dual output, native-space and template-space pathology, allows QNPtoVox to support both individualized anatomical analysis and population-level modeling within a single unified framework.

The value of a spatial standardization pipeline such as QNPtoVox increases substantially in the context of large, longitudinal cohort studies with extensive postmortem tissue repositories. Studies such as the Adult Changes in Thought (ACT) study and the Religious Orders Study and Memory and Aging Project (ROSMAP)^28^ have accumulated extensive collections of postmortem human brain tissue that have supported quantitative neuropathological analyses at scale. Among these resources, ACT is distinctive in its population-based design and the availability of ex vivo MRI acquired before tissue sectioning for many participants, providing a direct volumetric imaging scaffold for spatial alignment. We designed QNPtoVox to leverage such datasets by enabling cross-participant pooling of spatially registered pathology data, which is essential for examining subgroup differences, inter-individual heterogeneity, and spatial patterns of pathology that only emerge at large sample sizes.

Although ROSMAP and similar repositories do not systematically include ex vivo MRI, recent methodological advances offer viable alternatives for constructing three-dimensional anatomical reference volumes. In particular, techniques that computationally stitch serial gross coronal tissue slabs into volumetric brain reconstructions can provide an imaging scaffold that approximates native brain geometry^29^. These reconstructed volumes can serve as substitutes for ex vivo MRI in spatial registration workflows. QNPtoVox can incorporate slab-derived volumes as an anatomical reference for co-location and voxelization, thereby extending its applicability to large-scale brain tissue repositories that lack dedicated postmortem MRI. As these reconstruction methods mature, they offer a practical pathway for applying QNPtoVox to ROSMAP and other cohorts with extensive histopathology but limited volumetric imaging.

While we demonstrate QNPtoVox using AT8-immunostained tau pathology, the pipeline itself is stain-agnostic. QNPtoVox operates on quantitative pathology values and spatial coordinates extracted from histological images, independent of the biological marker being measured. Investigators can therefore apply the same workflow to other immunohistochemical stains, including markers of amyloid pathology, TDP-43, neuroinflammation, or vasculature. The pipeline can also accommodate quantitative markers of physiology, such as NeuN-based estimates of neuronal density. This flexibility supports the generation of spatially standardized voxel-level representations for diverse neuropathological features and enables integrative, multi-marker analyses within a unified anatomical framework.

Importantly, QNPtoVox is not limited to Alzheimer’s disease. The pipeline provides a generalizable spatial transformation framework that can be applied to quantitative neuropathology from other neurodegenerative disorders, including Lewy body disease, frontotemporal lobar degeneration, vascular cognitive impairment, and mixed pathologies. Any disease context in which pathology is assessed quantitatively in histological sections and benefits from spatial modeling can leverage QNPtoVox. This disease-agnostic design positions the pipeline as a broadly applicable method for integrating histopathology with neuroimaging-derived anatomical context across neurodegenerative research domains.

QNPtoVox is also quantitative pathology software agnostic. Although this implementation uses Halo for tile-level extraction, the pipeline does not depend on Halo-specific algorithms or proprietary data structures. Any digital pathology platform capable of exporting quantitative measurements together with spatial geometry can provide valid inputs. Open-source platforms such as QuPath support quantitative analysis with spatially explicit outputs that can be adapted to the QNPtoVox input specification. By decoupling spatial transformation from the choice of pathology software, QNPtoVox reduces vendor dependence and increases accessibility while allowing laboratories to retain their existing quantification workflows.

From a practical standpoint, QNPtoVox balances methodological rigor with feasibility. Assuming a standardized postmortem workup, quantitative pathology extraction (Step 1) typically requires approximately 30–60 minutes per histological slide, depending on staining quality and quality-control adjustments. Spatial transformation and registration (Step 2) require approximately 10 hours per participant’s brain. Nonlinear registration using ANTs^14^ constitutes the primary computational bottleneck, accounting for roughly 6–8 hours of runtime. However, users can parallelize this step and accelerate it on higher-end workstations or compute clusters. Expert-guided co-location introduces a limited manual component but remains efficient; identifying the appropriate coronal plane and in-plane offsets, combined with virtual slice review, typically requires no more than 30 minutes per participant. All remaining steps, including coordinate transformation, voxelization, smoothing, and voxel extraction, execute automatically and complete within minutes. These characteristics support application of QNPtoVox to moderately large cohorts without requiring specialized hardware.

Overall, QNPtoVox provides a reproducible and modular method for transforming quantitative neuropathology into voxel-aligned representations compatible with modern neuroimaging and spatial modeling frameworks. Rather than introducing a new biological metric or analytical model, the pipeline addresses a methodological gap that has historically separated histological precision from anatomical context. By enabling spatial standardization, cross-participant aggregation, and multimodal integration, QNPtoVox unlocks existing and future quantitative neuropathology datasets for scalable computational analysis. As digital pathology resources and three-dimensional tissue reconstruction methods continue to advance, QNPtoVox offers a flexible foundation for integrating quantitative neuropathology into the broader neuroimaging ecosystem across diseases, cohorts, and analytical paradigms.

## References

1. Mukhopadhyay S, Feldman MD, Abels E, et al. Whole Slide Imaging Versus Microscopy for Primary Diagnosis in Surgical Pathology: A Multicenter Blinded Randomized Noninferiority Study of 1992 Cases (Pivotal Study). The American Journal of Surgical Pathology. 2018;42(1):39. doi:10.1097/PAS.0000000000000948

2. HALO | Quantitative Image Analysis for Pathology - Indica Labs. Accessed August 2, 2025. https://indicalab.com/halo/

3. Bankhead P, Loughrey MB, Fernández JA, et al. QuPath: Open source software for digital pathology image analysis. Sci Rep. 2017;7(1):16878. doi:10.1038/s41598-017-17204-5

4. Braak H, Braak E. Demonstration of Amyloid Deposits and Neurofibrillary Changes in Whole Brain Sections. Brain Pathology. 1991;1(3):213–216. doi:10.1111/j.1750-3639.1991.tb00661.x

5. Braak H, Braak E. Staging of alzheimer’s disease-related neurofibrillary changes. Neurobiology of Aging. 1995;16(3):271–278. doi:10.1016/0197-4580(95)00021-6

6. Moms JC, Heyman A, Mohs RC, et al. The Consortium to Establish a Registry for Alzheimer’s Disease (CERAD). Part I. Clinical and neuropsychological assessment of Alzheimer’s disease. Neurology. 1989;39(9):1159–1159.

7. Thal DR, Rüb U, Orantes M, Braak H. Phases of A beta-deposition in the human brain and its relevance for the development of AD. Neurology. 2002;58(12):1791–1800. doi:10.1212/wnl.58.12.1791

8. Montine TJ, Phelps CH, Beach TG, et al. National Institute on Aging-Alzheimer’s Association guidelines for the neuropathologic assessment of Alzheimer’s disease: a practical approach. Acta Neuropathol. 2012;123(1):1–11. doi:10.1007/s00401-011-0910-3

9. Webster JM, Grabowski TJ, Madhyastha TM, Gibbons LE, Keene CD, Latimer CS. Leveraging Neuroimaging Tools to Assess Precision and Accuracy in an Alzheimer’s Disease Neuropathologic Sampling Protocol. Frontiers in Neuroscience. 2021;15:1061. doi:10.3389/FNINS.2021.693242/BIBTEX

10. Fonov V, Evans A, McKinstry R, Almli C, Collins D. Unbiased nonlinear average age-appropriate brain templates from birth to adulthood. NeuroImage. 2009;47:S102. doi:10.1016/S1053-8119(09)70884-5

11. Fischl B. FreeSurfer. NeuroImage. 2012;62(2):774-781. doi:10.1016/j.neuroimage.2012.01.021

12. Jenkinson M, Beckmann CF, Behrens TEJ, Woolrich MW, Smith SM. FSL. NeuroImage. 2012;62(2):782-790. doi:10.1016/j.neuroimage.2011.09.015

13. Penny WD, Friston KJ, Ashburner JT, Kiebel SJ, Nichols TE. Statistical Parametric Mapping: The Analysis of Functional Brain Images. Elsevier; 2011. Accessed January 6, 2026. https://books.google.com/books?hl=en&lr=&id=G_qdEsDlkp0C&oi=fnd&pg=PP1&dq=Statistical+Parametric+Mapping:+The+Analysis+of+Functional+Brain+Images.+Academic+Press.&ots=Xo2NFtY6WC&sig=FxYyVS2GlVZwElTlFbOueHnbISE

14. Avants BB, Tustison N, Song G. Advanced normalization tools (ANTS). Insight j. 2009;2(365):1–35.

15. Latimer CS, Melief EJ, Ariza-Torres J, et al. Protocol for the Systematic Fixation, Circuit-Based Sampling, and Qualitative and Quantitative Neuropathological Analysis of Human Brain Tissue. In: Chun J, ed. Alzheimer’s Disease: Methods and Protocols. Springer US; 2023:3–30. doi:10.1007/978-1-0716-2655-9_1

16. Adult Changes in Thought (ACT) Study. ACT Study Home. https://actagingresearch.org/index.php?cID=308#data-sharing-policies

17. Montine TJ, Monsell SE, Beach TG, et al. Multisite assessment of NIA-AA guidelines for the neuropathologic evaluation of Alzheimer’s disease. Alzheimers Dement. 2016;12(2):164–169. doi:10.1016/j.jalz.2015.07.492

18. Welcome to Python.org. Python.org. December 16, 2025. Accessed January 6, 2026. https://www.python.org/

19. Array programming with NumPy | Nature. Accessed January 6, 2026. https://www.nature.com/articles/s41586-020-2649-2

20. McKinney W. Data structures for statistical computing in Python. scipy. 2010;445(1):51-56.

21. SciPy 1.0: fundamental algorithms for scientific computing in Python | Nature Methods. Accessed August 5, 2025. https://www.nature.com/articles/s41592-019-0686-2

22. Brett M, Markiewicz CJ, Hanke M, et al. nipy/nibabel: 5.3.1. Published online October 15, 2024. doi:10.5281/zenodo.13936989

23. R: The R Project for Statistical Computing. Accessed January 6, 2026. https://www.r-project.org/

24. Clayden [cre J, aut, Cox B, et al. RNifti: Fast R and C++ Access to NIfTI Images. Published online February 22, 2025. Accessed January 6, 2026. https://cran.r-project.org/web/packages/RNifti/index.html

25. Urbanek S. png: Read and write PNG images. Published online November 29, 2022. Accessed January 6, 2026. https://cran.r-project.org/web/packages/png/index.html

26. McCarthy P. FSLeyes. Published online May 29, 2025. doi:10.5281/zenodo.15542963

27. Horn A. MNI T1 6thGen NLIN to MNI 2009b NLIN ANTs transform. Published online 2016:208235992 Bytes. doi:10.6084/M9.FIGSHARE.3502238.V1

28. Bennett DA, Buchman AS, Boyle PA, Barnes LL, Wilson RS, Schneider JA. Religious Orders Study and Rush Memory and Aging Project. J Alzheimers Dis. 2018;64(Suppl 1):S161-S189. doi:10.3233/JAD-179939

29. Gazula H, Tregidgo HF, Billot B, et al. Machine learning of dissection photographs and surface scanning for quantitative 3D neuropathology. Zhou JH, Makin TR, eds. eLife. 2024;12:RP91398. doi:10.7554/eLife.91398

